# Pulse train gating to improve signal generation for *in vivo* two-photon fluorescence microscopy

**DOI:** 10.1101/2023.04.03.535393

**Authors:** Shaun A. Engelmann, Alankrit Tomar, Aaron L. Woods, Andrew K. Dunn

## Abstract

**Significance:** Two-photon microscopy is used routinely for *in vivo* imaging of neural and vascular structure and function in rodents with a high resolution. Image quality, however, often degrades in deeper portions of the cerebral cortex. Strategies to improve deep imaging are therefore needed. We introduce such a strategy using gates of high repetition rate ultrafast pulse trains to increase signal level.

**Aim:** We investigate how signal generation, signal-to-noise ratio (SNR), and signal-to-background ratio (SBR) improve with pulse gating while imaging *in vivo* mouse cerebral vasculature.

**Approach:** An electro-optic modulator is used with a high-power (6 W) 80 MHz repetition rate ytterbium fiber amplifier to create gates of pulses at a 1 MHz repetition rate. We first measure signal generation from a Texas Red solution in a cuvette to characterize the system with no gating and at a 50%, 25%, and 12.5% duty cycle. We then compare signal generation, SNR, and SBR when imaging Texas Red-labeled vasculature using these conditions.

**Results:** We find up to a 6.73-fold increase in fluorescent signal from a cuvette when using a 12.5% duty cycle pulse gating excitation pattern as opposed to a constant 80 MHz pulse train. We verify similar increases for *in vivo* imaging to that observed in cuvette testing. For deep imaging we find pulse gating to result in a 2.95-fold increase in SNR and a 1.37-fold increase in SBR on average when imaging mouse cortical vasculature at depths ranging from 950 μm to 1050 μm.

**Conclusions:** We demonstrate that a pulse gating strategy can either be used to limit heating when imaging superficial brain regions or used to increase signal generation in deep regions. These findings should encourage others to adopt similar pulse gating excitation schemes for imaging neural structure through two-photon microscopy.

## 1 Introduction

Two-photon microscopy is routinely used to image *in vivo* neural structure throughout the cerebral cortex in rodents at a micrometer resolution^1-4^. Ultrafast lasers are focused using high numerical aperture objectives and raster scanned across an imaging plane to excite fluorophores which label anatomical structure. A natural limit to imaging depth arises, however, once the background fluorescence generated away of the focus approaches the level of fluorescence produced at the imaging plane^5^ or when the attenuation of excitation and emission light due to tissue scattering reduces the image signal to a point where it is no longer detectable. The unavoidable reduction in signal-to-background ratio (SBR) with depth critically inhibits the ability to accurately image deep anatomical structure. The issue is further exacerbated by the inherent noise that materializes during signal detection, which makes a high signal-to-noise ratio (SNR) crucial as well.

Many strategies exist to improve deep imaging in tissue. One is to use a longer excitation wavelength that is less susceptible to scattering as it travels through tissue, leading to a higher percentage of photons at the focal point for a given average power, ultimately increasing signal^6-9^. Long excitation wavelengths also enable three-photon excitation with select fluorophores, which significantly reduces the amount of background florescence generated relative to two-photon imaging schemes^10,11^. Long wavelength and three-photon imaging both extend the theoretically possible imaging depth. A more trivial solution to improve imaging at depth is to simply use higher excitation power. This does not increase the theoretical SBR-imposed depth limit, but rather it increases SBR in all regions above this point ultimately facilitating imaging to the theoretical limit. However, this approach is ultimately limited by the need to limit average power levels to avoid excessive tissue heating from light absorption^12,13^.

Since heating is a function of average power and signal is related to the square of peak power (or cube in three-photon excitation), it can be useful to reduce the pulse repetition rate and use higher pulse energies while remaining below thermal damage thresholds. For this reason, low repetition rate (∼1 MHz) sources are often used for multiphoton imaging^14,15^. These sources are commercially available but are complex and very expensive. An alternative approach is to reduce pulse delivery from high repetition rate, high average power ultrafast lasers through pulse picking or pulse gating. There has not yet been a thorough quantification of the improvements brought forth by pulse reduction while holding pulse characteristics (pulse width, pulse spectrum) constant for *in vivo* imaging. Our aim with this work is to provide such a comparison using a high power, relatively inexpensive, high repetition rate (80 MHz) excitation source and a pulse gating system. Past work with similar strategies has either been limited to *ex vivo* imaging^16,17^, or performed with an excitation system where pulse characteristics varied with repetition rate, complicating interpretation of the results^18^. What is presented here should encourage adoption of similar pulse train gating schemes for *in vivo* deep brain imaging.

## 2 Methods

### 2.1 Ultrafast Pulse Gating System

The complete excitation and microscopy setup is shown in Fig. 1A. A custom ytterbium fiber amplifier served as the excitation laser for this work^19,20^. The amplifier was constructed using 6 m of double-clad polarization-maintaining ytterbium-doped fiber. It is seeded by an 80 MHz commercial oscillator with a 100 mW average power (NKT Photonics, Origami 10-80) and pumped by a 30 W, 915 nm laser diode. Amplifier settings were selected to produce a beam with an average power of 6 W after pulse compression and dispersion compensation, which was accomplished using a pair of transmission gratings. Pulses are centered around λ=1060 nm (Fig. 1B), and *in situ* autocorrelation indicates a pulse width of 110 fs in the imaging plane assuming a sech^2^ pulse shape (Fig. 1C). Complete details of the Yb amplifier can be found in previous publications^19,20^.

**Fig. 1.**
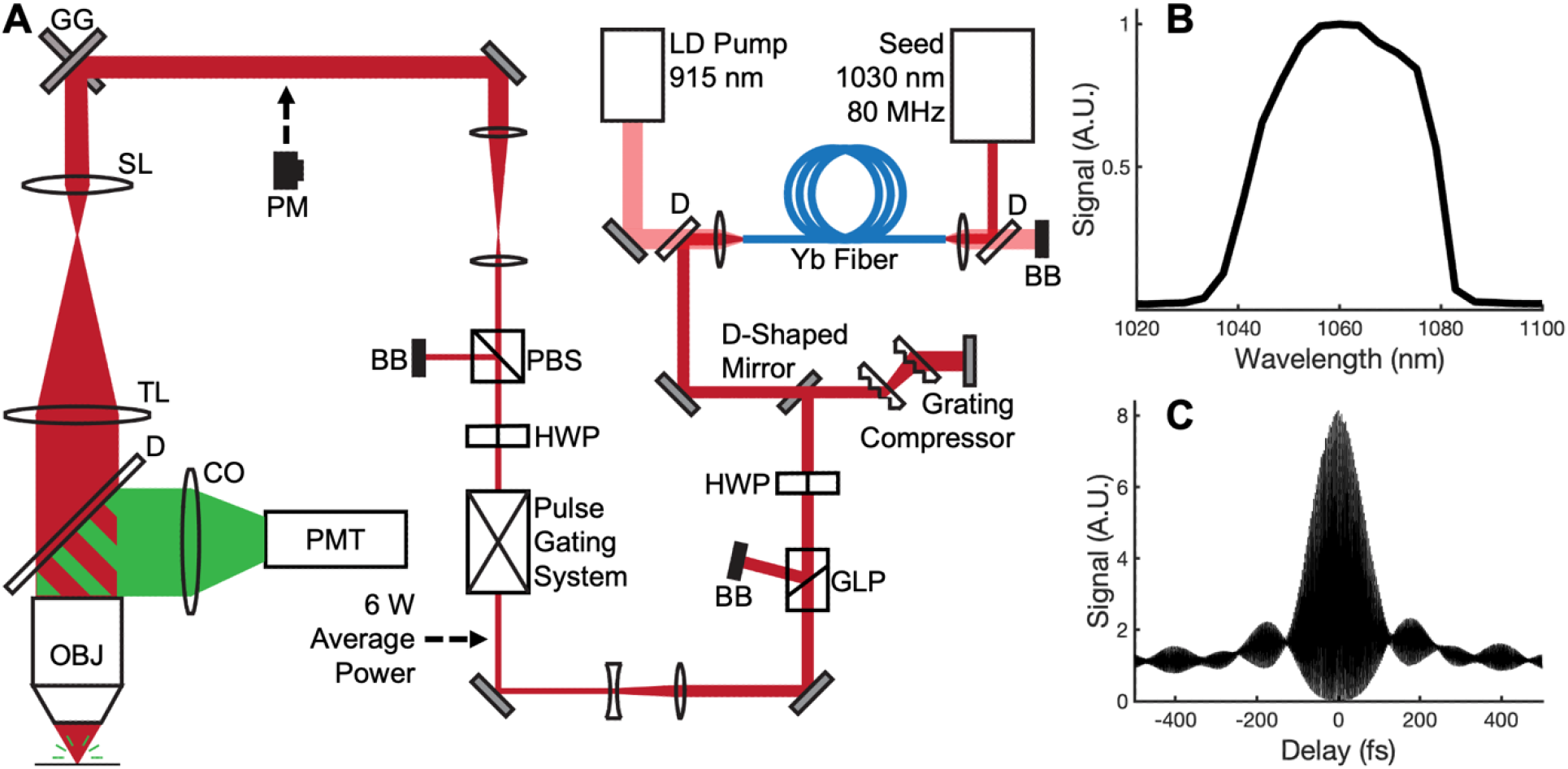
Schematic and characterization of the imaging system. (A) Schematic of the fiber amplifier, pulse gating system, and two-photon microscope. (B) Fiber amplifier spectrum. (C) *In situ* fiber amplifier autocorrelation measured through the objective. The pulse width is approximately 110 fs assuming a sech^2^ shape. GG - galvo-galvo scanners, SL – scan lens, TL – tube lens, D – dichroic, OBJ – objective, CO-collection optics, PMT – photomultiplier tube, PM – power meter, BB-beam block, PBS – polarizing beam splitter, HWP – half-waveplate, GLP – Glan-laser polarizer, LD – laser diode.

Following compression, the beam is telescoped down to a diameter of ∼1.5 mm and sent through an electro-optic modulator (EOM, Conoptics, 360-80) for pulse gating. The extinction ratio for rejected pulses is approximately 100:1, and the throughput is ∼85% for transmitted light. The EOM is controlled by a driver (Conoptics, Model 25D) which either passes or rejects light based on a signal from a digital delay generator (Stanford Research System, DG645) triggered off the amplifier seed. The EOM control pulses were designed to pass gates of excitation pulses at a 1 MHz repetition rate as shown in Fig. 2. Gate duration was set to achieve a duty cycle of either 50%, 25%, or 12.5%. A 25% duty cycle, for example, would cyclically pass 20 pulses and then reject 60 from the 80 MHz fiber amplifier pulse train. This has a similar effect to reducing the original 80 MHz repetition rate to 20 MHz, which was not possible given the timing limitations of the gating equipment used.

**Fig. 2.**
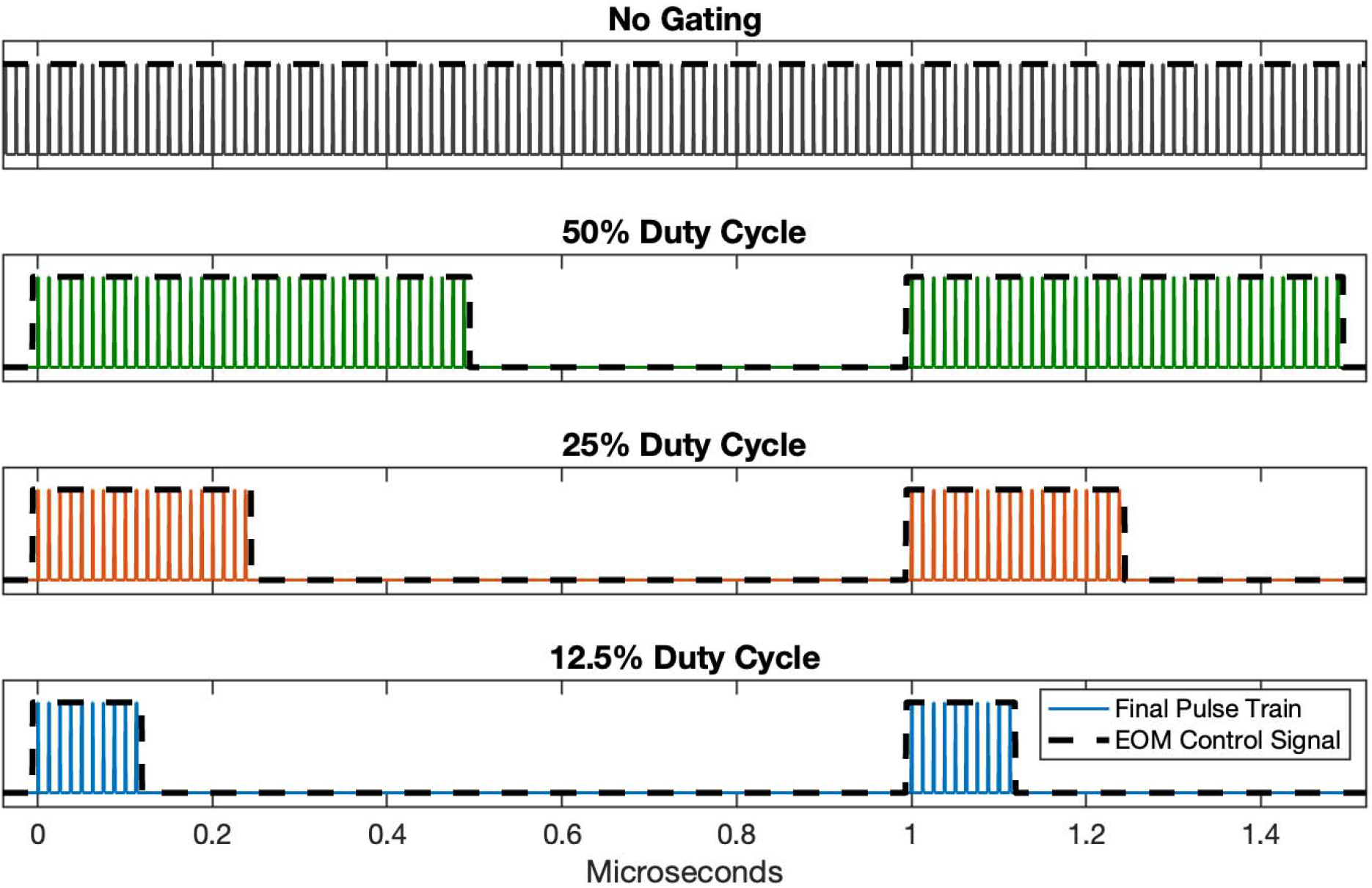
Ideal pulse trains used for imaging. There is a “no gating” condition where the EOM was programmed to transmit all pulses in the 80 MHz fiber amplifier pulse train, as well as 50%, 25%, and 12.5% duty cycle conditions where the EOM transmitted pulse trains in gates at a 1 MHz repetition rate.

### 2.2 Multiphoton Microscope

A half-waveplate and polarizing beam splitter set the excitation power sent to the microscope following pulse gating. The beam was expanded to fill the microscope objective back aperture, which was either a 25X objective (Olympus XLPLN25XSVMP2, 1.0NA) for *in vivo* imaging or a 10X objective (Nikon CFIPlan10X, 0.25NA) for cuvette experimentation. The excitation beam was scanned using a pair of galvanometer mirrors (Thorlabs, QS7XY-AG) conjugated the objective back aperture using a scan lens (Thorlabs, SL50-2P2) and Plössl tube lens (Thorlabs, AC508-400-C) combination. Backscattered fluorescent signal was directed to a photomultiplier tube (PMT, Hamamatsu, H10770PB-50) with a 775 nm cutoff dichroic (Semrock, FF775-Di01), and was further filtered with a 609/181 bandpass filter (Semrock, FF01-609/181-25) which immediately preceded the PMT photocathode. Imaging was controlled using a custom LabVIEW software. While it may be necessary to synchronize EOM gating with DAQ sampling in some situations, this was not done for this work given our low sampling rate (160 kHz) relative to the gating frequency (1 MHz).

### 2.3 Animal Protocols

All animal work was approved of by the University of Texas at Austin Institutional Animal Care and Use Committee. Adult female C57BL/6 mice were fit with cranial windows in a similar manner to that described in a previous publication^21^. Carprofen (10 mg/kg, subcutaneous) and dexamethasone sodium phosphate (2mg/kg, subcutaneous) were administered to control inflammation during the procedure, and mice were allowed to heal for at least two weeks prior to imaging. Temperature was maintained using a heating pad during both surgeries and imaging, where mice were anesthetized with isoflurane (2.5% induction, 1.5% maintenance). In all *in vivo* sessions, vasculature was labeled through 100 μL retro-orbital injections of 5% w/v 70 kDa dextran-conjugated Texas Red diluted in physiological saline. Excitation power was limited to 100 mW measured at the brain surface to avoid thermal damage.

## 3 Results

### 3.1 Signal Generation in a Cuvette as a Function of Excitation Duty Cycle

The pulse gating system was set to either transmit the complete 80 MHz pulse train without gating, or to transmit pulse trains with a 12.5%, 25%, or 50% duty cycle at a 1 MHz repetition rate. Fluorescent signal generated within a cuvette containing a 39 μM Texas Red solution was measured for all gating conditions. Average excitation power for each state was 5 mW at minimum (measured before the microscope optics at the position shown in Fig. 1A) and increased by factors of √2 until saturation was observed. Plots of the signal level measured for each excitation condition are shown in Fig. 3. When plotting fluorescence versus the excitation power on a log-log scale, linear fits reveal a slope of 2 for all conditions as expected for two-photon excitation. For the 50%, 25%, and 12.5% duty cycles we see a 2.02, 3.97, and 6.73-fold average increase in signal respectively relative to the no gating condition.

**Fig. 3.**
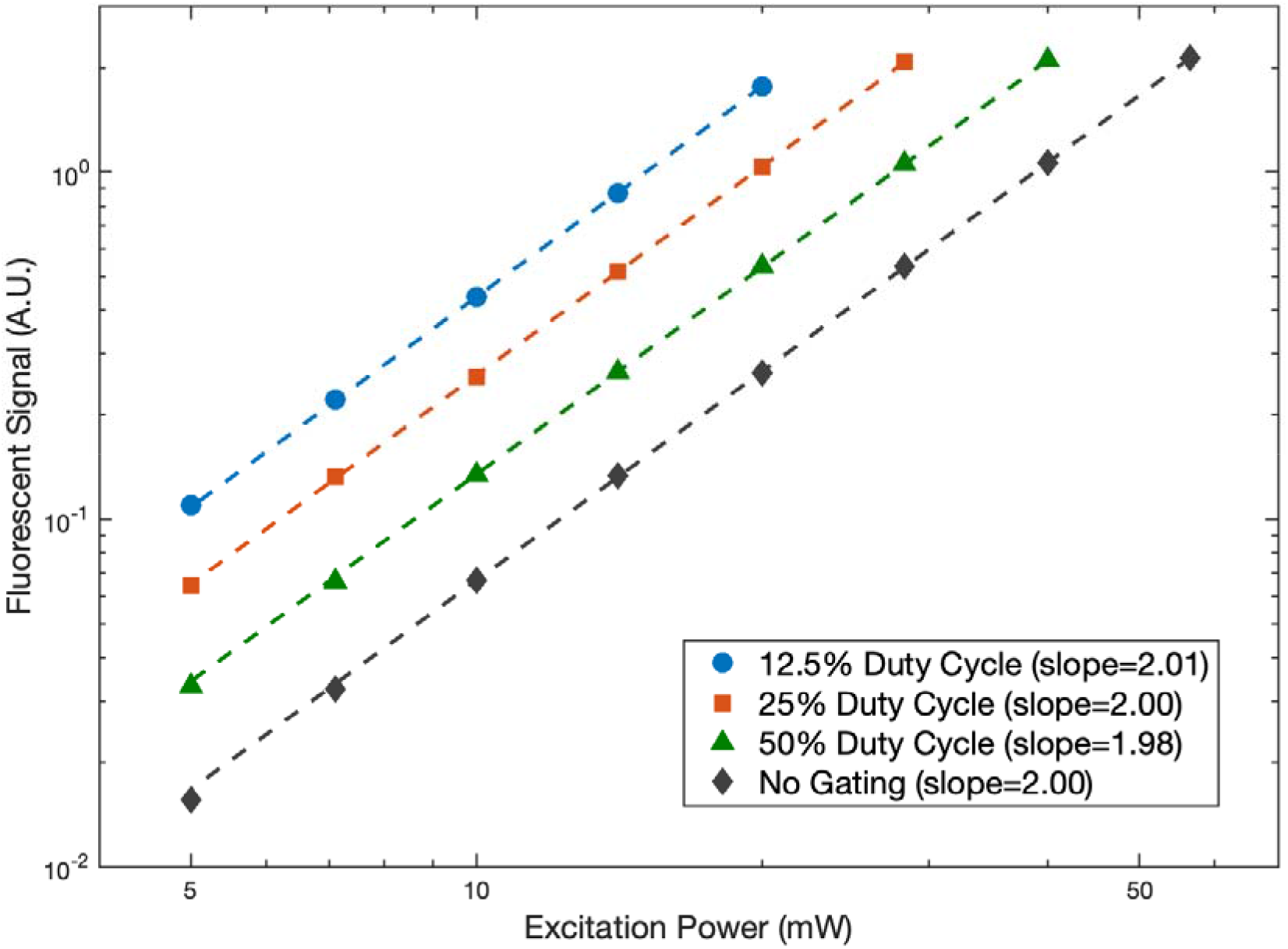
Fluorescent signal generated with different excitation duty cycles. Power was varied from 5 to a maximum of 56.6 mW measured before the microscope optics at the position shown in Fig. 1A. The resulting fluorescence measurements are plotted with the pre-microscope power on a log-log scale. Linear fits were performed for each duty cycle (dashed lines) and the resulting slopes are included in the plot legend.

### 3.2 In Vivo Signal Generation Comparison between Duty Cycles

For *in vivo* imaging, the pulse gating system was set to transmit pulse trains with either a 12.5%, 25%, or 50% duty cycle at a 1MHz repetition rate. The 80 MHz ungated pulse train was not used here as the system required an elongated period for stabilization following a switch to this condition due to thermal effects within the EOM crystal (Fig. S1). First, a 12.5% duty cycle pulse train was used to image a 3-slice stack (3 μm axial step size, 8 frames averaged) where power level was set to avoid saturation. Stacks were then reacquired with this same duty cycle using 75% and 50% of the original power. This was followed by the acquisition of two final stacks both at the original power using a 25% and then a 50% duty cycle. This process was performed twice at approximate depths of 250 μm and 500 μm.

Following acquisition, 1-pixel-radius median filters were applied to each image and maximum intensity projections were created to ultimately use for analysis (Fig. 4A). 5 pixel-thick, 80 pixel-long line profiles were created for the same 3 vessels in each projection at each depth. Figure 4B shows example profiles for the vessels indicated in Fig. 4A. Profiles are shown for the additional vessels in Fig. S2 and Fig. S3. SBR was determined with the profiles by dividing the peak value by the background, which was the average of the first 15 and last 15 pixels. Relative to the 25% and 50% duty cycles, the 12.5% duty cycle condition resulted in peaks that were on average 2.02 ± 0.15 and 3.59 ± 0.37 times greater in magnitude respectively (± indicating standard error). Relative to the 75% and 50% average power conditions, the full power 12.5% duty cycle condition resulted in peaks that were on average 1.84 ± 0.15 and 4.42 ± 0.23 greater in magnitude respectively. No major differences in relative peak signal generation between the excitation conditions are observed when comparing these two depths.

**Fig. 4.**
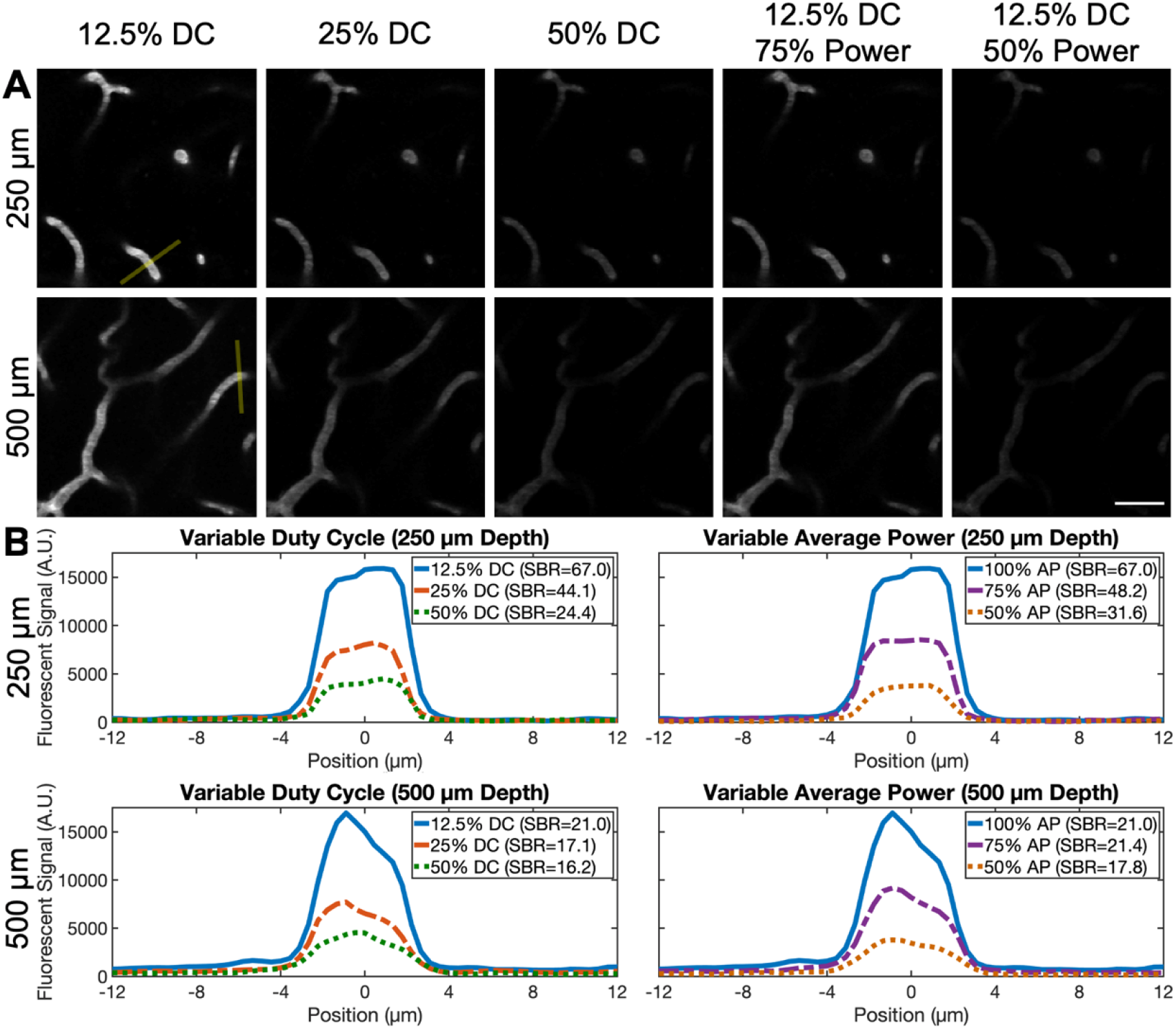
*In vivo* comparison of pulse delivery conditions. (A) Images of Texas Red-labeled vasculature acquired using varying excitation powers and duty cycles at approximately 250 μm (top) and 500 μm (bottom) deep. Each one is a 3-image maximum intensity projection (3 μm axial step size). 5 pixel-thick lines for the vessels indicated in (A) were used to create the profiles in (B) and to calculate SBR. Average power at the brain surface was held constant for the leftmost line profiles (7 mW for 250 μm, 31 mW for 500 μm) and duty cycle was held constant for the rightmost line profiles (12.5% for 250 μm and 500 μm) in (B). Scale bar is 25 μm.

### 3.3 Pulse Gating for Deep Imaging

To investigate pulse gating when imaging deep structure, the 12.5% duty cycle pulse train was directly compared to the no gating condition. The EOM was first set to transmit the complete 80 MHz pulse train, and Texas Red-labeled neurovasculature was imaged in 5 μm axial step sizes until vessels were no longer clearly observed. Both frame averaging and average excitation power were increased with depth until 925 μm was reached. Then from 925 μm to 1125 μm, 10 frames were averaged together for each acquired image and excitation power at the brain surface was maintained at 100 mW. The pulse gating system was then altered to operate at a 12.5% duty cycle, and EOM throughput was allowed to equalize for approximately 10 minutes as a thermal equilibrium was approached (Fig. S1). Despite the thermal differences, pulse width is the same for each gating condition (Fig. S4). Vasculature was then reimaged from 925 μm to 1125 μm using the same average power and frame averaging conditions as with the ungated pulse train. Maximum intensity projections through the resulting deep vascular stacks are displayed in Fig. 5. To quantify benefits of using a reduced pulse delivery, 3 representative vessels at 950, 1000, 1050, and 1100 μm were used to calculate both SNR and SBR (Fig. 6A). Similarly to Sec. 3.2, the images for analysis were projections created using the image at the indicated depth along with those at the preceding and following depths. SBR was calculated in the same manner previously described, while SNR was calculated using the formula (μ_peak_-μ_bkgr_)/σ_bkgr_, where the 5 greatest values were averaged for μ_peak_ and the 15 first and 15 last pixels were used for μ_bkgr_ and σ_bkgr_. A greater SNR and SBR are observed for all depths for the 12.5% duty cycle pulse train as shown in Fig. 6C and 6D. Table 1 lists the relative increase in these two metrics at each depth. The profiles for one vessel at each depth are shown in Fig. 6B, and profiles for the remaining vessels are shown in Fig. S5.

**Fig. 5.**
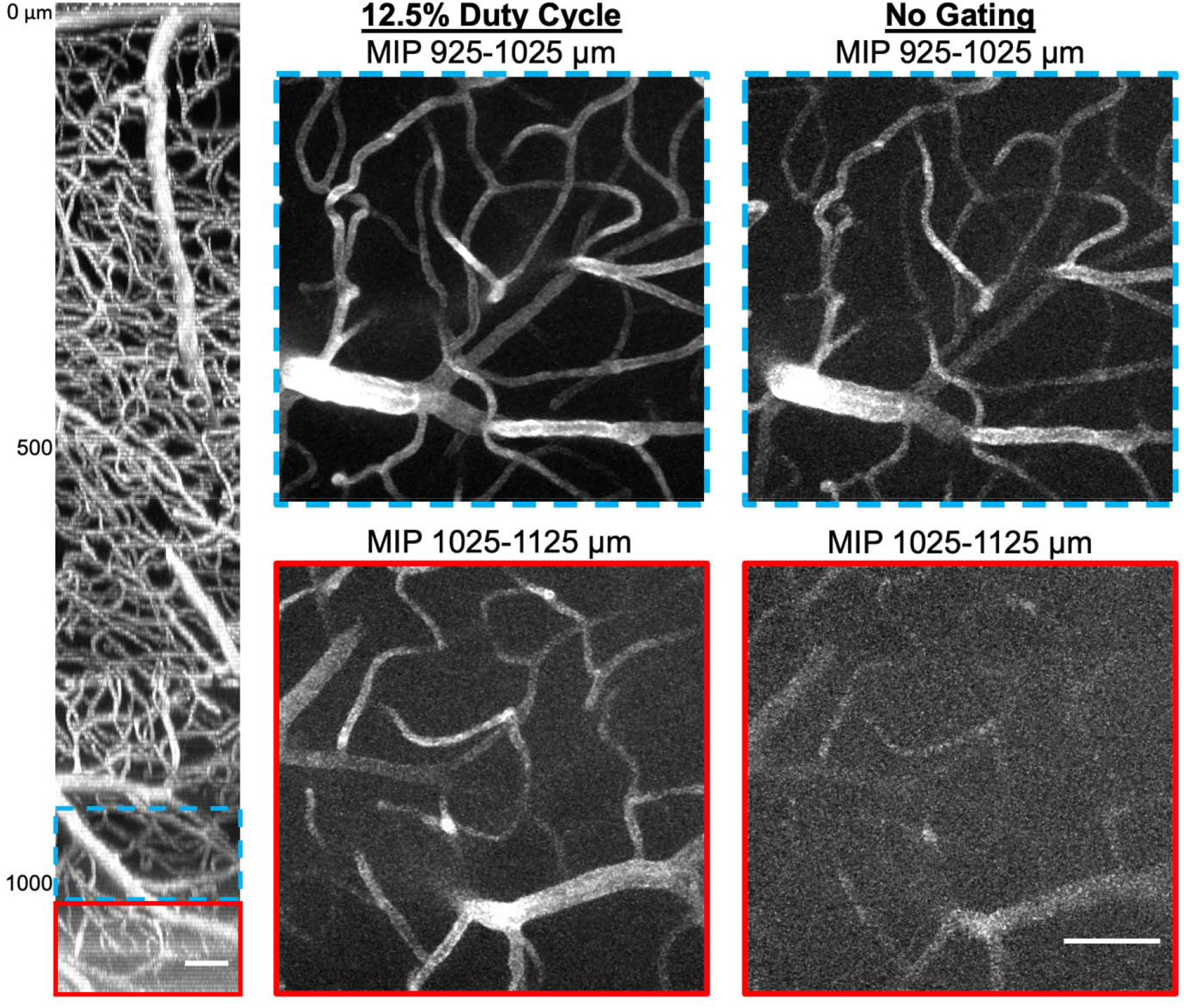
Deep imaging of Texas Red-labeled vasculature at a 12.5% duty cycle (middle) and without pulse gating (right). These maximum intensity projections (MIP) were recorded using a 100 mW average power measured at the brain surface. The side-projection (left) was imaged using <5mW at the surface and up to 100 mW in deep regions. Scale bars are 50 μm.

**Table 1.**
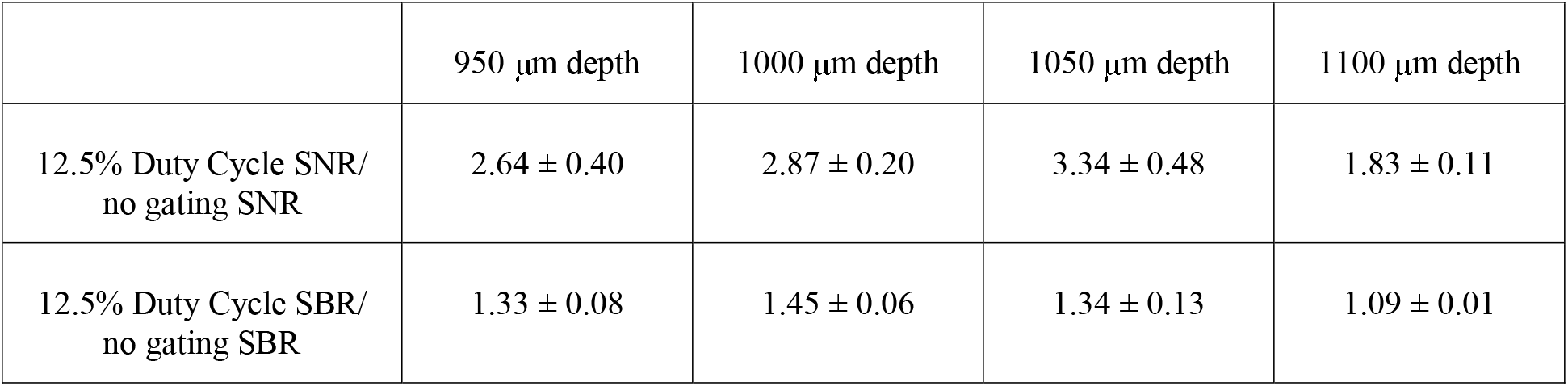
Relative signal-to-noise and signal-to-background ratio (SNR, SBR) using a 12.5% duty cycle pulse gating strategy compared to imaging with no gating. Values were calculated using the same 3 vessel profiles at each depth shown in Fig. 6. Values after the ± indicate standard error.

| | 950 $\mu\text{m}$ depth | 1000 $\mu\text{m}$ depth | 1050 $\mu\text{m}$ depth | 1100 $\mu\text{m}$ depth |
| --- | --- | --- | --- | --- |
| 12.5% Duty Cycle SNR/<br>no gating SNR | $2.64 \pm 0.40$ | $2.87 \pm 0.20$ | $3.34 \pm 0.48$ | $1.83 \pm 0.11$ |
| 12.5% Duty Cycle SBR/<br>no gating SBR | $1.33 \pm 0.08$ | $1.45 \pm 0.06$ | $1.34 \pm 0.13$ | $1.09 \pm 0.01$ |

**Fig. 6.**
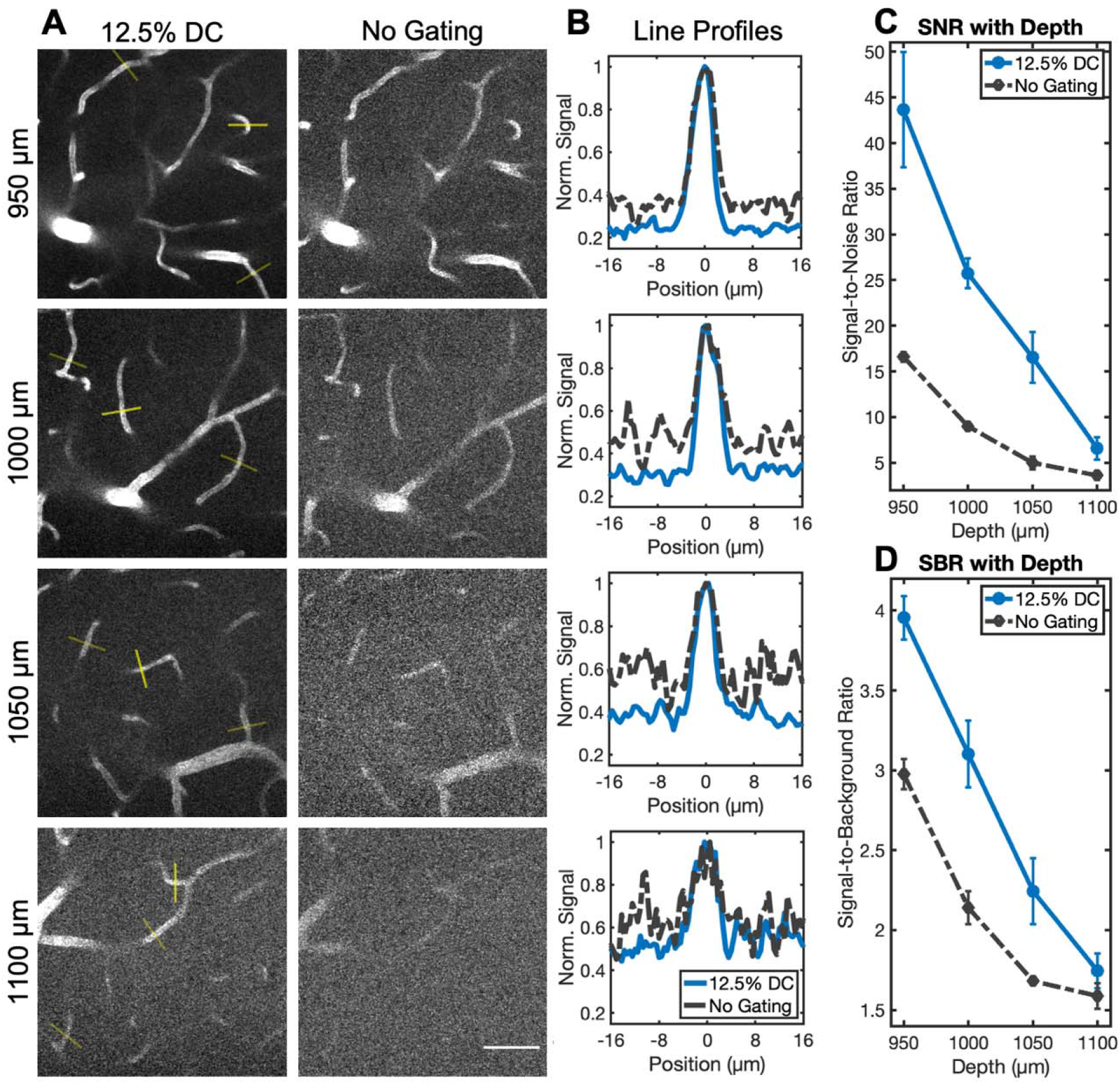
Deep imaging signal-to-noise ratio (SNR) and signal-to-background ratio (SBR) comparisons between the 12.5% duty cycle and the ungated conditions. (A) Images used for quantifying SNR and SBR. Each one is a 3-image maximum intensity projection centered around the indicated depth (5 μm axial step size). (B) Line profiles through the vessels with the brightest markers shown in (A). Each plot corresponds with the depth listed in the same row. (C) Average SNR with depth as determined from all vessels indicated in (A). (D). Average SBR with depth for all vessels in (A). Images are from the same stacks displayed in Fig. 5. Power at the brain surface was 100 mW for all images. Scale bar is 50 μm. Error bars represent standard error.

## 4 Discussion

To verify proper system behavior, it is helpful to calculate the theoretically expected enhancements in signal with pulse gating. Fluorescence intensity (*F*) for two-photon excitation is proportional to the square of pulse energy (*E*), *F* ∝ *E*^*2*^. When reducing the 80 MHz pulse train using a 12.5% duty cycle, we would expect an 8-fold increase in signal when imaging with the same average power as shown by comparing

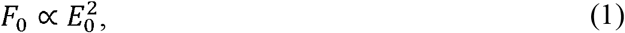

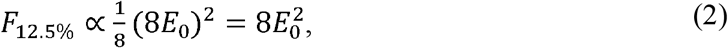

where *F*_*0*_ and *E*_*0*_ are the fluorescence intensity and pulse energy for the ungated pulse train, and *F*_*12*.*5%*_ is fluorescence intensity for 12.5% duty cycle gating. Here we assume that the pulse energy is increased by 8 since the average power remains constant, but the number of pulses is reduced by a factor of 8. Equation 2 assumes that the pulse characteristics are constant between the two different conditions (verified in Fig. S4). Equation 2, however, does not account for the imperfect extinction and transmission of the EOM. To get a more accurate pulse energy relationship for our system (1:100 extinction ratio, 85% transmission) the following must be set equal

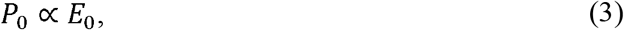

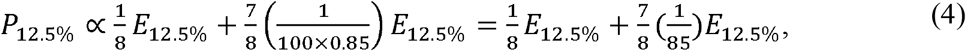

where *P*_*0*_ and *P*_*12*.*5%*_ are average powers. This yields a relationship of E_12.5%_ ≈ 7.39E_0_. Taking this, we can modify Eq. 2 as

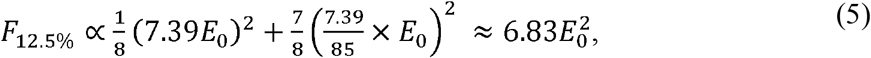

where we now expect 6.83-fold increase in signal magnitude for the 12.5% duty cycle as compared to the ungated condition (Eq. 2). Experimentally we found a similar increase of 6.73 when exciting Texas Red in solution, verifying that our pulse gating strategy improves signal generation as expected.

After completing this procedure for all evaluated duty cycles, we expect the 12.5% duty cycle to result in a 3.50-fold and 1.83-fold increase in signal as compared to the 50% and 25% duty cycles respectively. We found similar 3.33-fold and 1.69-fold increases respectively during cuvette testing, and 3.59-fold and 2.02-fold increases in peak signal during superficial vascular imaging. We have made use of this increase to demonstrate two useful applications of pulse gating for *in vivo* imaging. First, reducing the duty cycle allows imaging with lower average powers to limit tissue heating while maintaining high signal levels (demonstrated in Sec. 3.2). Second, images can be acquired with a reduced pulse delivery at the same average power to better resolve features. We use this strategy to improve deep vascular imaging in Sec. 3.3 where we observe consistent increases in both SBR and SNR. This can help researchers accurately identify vessels in images to better vectorize neurovascular networks^22,23^. Note that the SBR will approach 1 at the same depth for all gating conditions. This suggests that the relative benefit of pulse reduction will decrease with depth, which seems to be confirmed in Table 1 where the smallest improvement in SBR is seen at the deepest location analyzed (1100 μm). That said, many vessels that are nearly indistinguishable here when imaging with the ungated 80 MHz pulse train are resolved when switching over to the 12.5% duty cycle excitation pattern. This can in some part be attributed to the improved SNR offered alongside the raised SBR.

The ability to image with pulse gates at such a low duty cycle and still have acceptable average powers for deep two-photon microscopy is enabled by the relatively high average power of our custom fiber amplifier. The entire excitation system is relatively cost-efficient^19-21^ compared to commercial alternatives that provide similar pulse energies. The imaging strategy can be modified to further limit average power by synchronizing pulse delivery to only occur while scanning discrete regions of interest in a similar manner to Li et al^24^. While the system used in this study has an advantage over ours in that it can excite fluorophores through either three-photon or two-photon excitation, ours has a relatively simpler construction and is therefore more accessible. Our two-photon excitation wavelength is also slightly longer (1060 nm as compared to 920 nm), which further assists with deep imaging when paired with appropriate fluorophores. This work should ultimately encourage other groups, including those with limited optics expertise, to adopt similar pulse gating systems for deep imaging applications where potential thermal damage is a concern.

## 5 Conclusion

In this work we use an EOM-based pulse gating system to produce up to a 6.73-fold increase in two-photon signal generation while maintaining excitation power. We additionally demonstrate that pulse gating improves *in vivo* neurovascular two-photon imaging by raising both SBR and SNR. We demonstrate an average 1.37-fold increase in SBR and a 2.95-fold increase in SNR when imaging from 950 μm to 1050 μm in depth with 12.5% duty cycle pulse train gating as opposed to using a constant 80 MHz pulse train with the same average power. Around a 1100 μm depth, the pulse gating helps resolve vessels that are otherwise indistinguishable for the frame averaging conditions used. The results presented here should encourage use of pulse gating as a viable method to either mitigate thermal damage through reducing average excitation power, or to improve SNR and SBR through allowing heightened excitation pulse energies.

## Acknowledgments

We acknowledge funding from the National Institutes of Health (NS108484, T32EB007507) and the UT Austin Portugal Program.

## Code, Data, and Materials Availability

Raw images and data are not publicly available at this time but may be obtained from the authors upon reasonable request

## References

1. F. Helmchen, and W. Denk, “Deep tissue two-photon microscopy,” Nature methods 2(12), 932–940 (2005).

2. K. Svoboda, and R. Yasuda, “Principles of two-photon excitation microscopy and its applications to neuroscience,” Neuron 50(6), 823–839 (2006).

3. A. Y. Shih, J. D. Driscoll, P. J. Drew, N. Nishimura, C. B. Schaffer, and D. Kleinfeld, “Two-photon microscopy as a tool to study blood flow and neurovascular coupling in the rodent brain,” Journal of Cerebral Blood Flow & Metabolism 32(7), 1277–1309 (2012).

4. C. Grienberger, A. Giovannucci, W. Zeiger, and C. Portera-Cailliau, “Two-photon calcium imaging of neuronal activity,” Nature Reviews Methods Primers 2(1), 67 (2022).

5. P. Theer, and W. Denk, “On the fundamental imaging-depth limit in two-photon microscopy,” JOSA A 23(12), 3139–3149 (2006).

6. D. Kobat, M. E. Durst, N. Nishimura, A. W. Wong, C. B. Schaffer, and C. Xu, “Deep tissue multiphoton microscopy using longer wavelength excitation,” Optics express 17(16), 13354–13364 (2009).

7. D. Kobat, N. G. Horton, and C. Xu, “In vivo two-photon microscopy to 1.6-mm depth in mouse cortex,” Journal of biomedical optics 16(10), 106014–106014 (2011).

8. D. R. Miller, A. M. Hassan, J. W. Jarrett, F. A. Medina, E. P. Perillo, K. Hagan, S.M.S. Kazmi et al. “In vivo multiphoton imaging of a diverse array of fluorophores to investigate deep neurovascular structure,” Biomedical optics express 8(7), 3470–3481 (2017).

9. D. R. Miller, J. W. Jarrett, A. M. Hassan, and A. K. Dunn, “Deep tissue imaging with multiphoton fluorescence microscopy.” Current opinion in biomedical engineering 4, 32–39 (2017).

10. N. G. Horton, K. Wang, D. Kobat, C. G. Clark, F. W. Wise, C. B. Schaffer, and C. Xu, “In vivo three-photon microscopy of subcortical structures within an intact mouse brain,” Nature photonics 7(3), 205–209 (2013).

11. T. Wang, and C. Xu, “Three-photon neuronal imaging in deep mouse brain,” Optica 7(8), 947–960 (2020).

12. T. Wang, C. Wu, D. G. Ouzounov, W. Gu, F. Xia, M. Kim, X. Yang, M. R. Warden, and C. Xu, “Quantitative analysis of 1300-nm three-photon calcium imaging in the mouse brain,” Elife 9, e53205 (2020).

13. K. Podgorski, and G. Ranganathan, “Brain heating induced by near-infrared lasers during multiphoton microscopy,” Journal of neurophysiology 116(3), 1012–1023 (2016).

14. E. Beaurepaire, M. Oheim, and J. Mertz, “Ultra-deep two-photon fluorescence excitation in turbid media,” Optics Communications 188(1-4), 25–29 (2001).

15. P. Theer, M. T. Hasan, and W. Denk, “Two-photon imaging to a depth of 1000 µm in living brains by use of a Ti:Al2O3 regenerative amplifier,” Optics letters 28(12), 1022–1024 (2003).

16. D. Stachowiak, J. Bogusławski, A. Głuszek, Z. Łaszczych, M. Wojtkowski, and G. Soboń, “Frequency-doubled femtosecond Er-doped fiber laser for two-photon excited fluorescence imaging,” Biomedical Optics Express 11(8), 4431–4442 (2020).

17. V. Gautam, J. Drury, J. M. C. Choy, C. Stricker, H-A. Bachor, and V. R. Daria, “Improved two-photon imaging of living neurons in brain tissue through temporal gating,” Biomedical optics express 6(10), 4027–4036 (2015).

18. K. Charan, B. Li, M. Wang, C. P. Lin, and C. Xu, “Fiber-based tunable repetition rate source for deep tissue two-photon fluorescence microscopy,” Biomedical optics express 9(5), 2304–2311 (2018).

19. S. A. Engelmann, A. Zhou, A. M. Hassan, M. R. Williamson, J. W. Jarrett, E. P. Perillo, A. Tomar, D. J. Spence, T. A. Jones, and A. K. Dunn, “Diamond Raman laser and Yb fiber amplifier for in vivo multiphoton fluorescence microscopy,” Biomedical Optics Express 13(4), 1888–1898 (2022):.

20. E. P. Perillo, J. W. Jarrett, Y-L. Liu, A. Hassan, D. C. Fernée, J. R. Goldak, A. Bonteanu, D. J. Spence, H-C. Yeh, and A. K. Dunn, “Two-color multiphoton in vivo imaging with a femtosecond diamond Raman laser,” Light: Science & Applications 6(11), e17095–e17095 (2017).

21. E. P. Perillo, J. E. McCracken, D. C. Fernée, J. R. Goldak, F. A. Medina, D. R. Miller, H-C. Yeh, and K. Dunn, “Deep in vivo two-photon microscopy with a low cost custom built mode-locked 1060 nm fiber laser,” Biomedical optics express 7(2), 324–334 (2016).

22. A. Zhou, S. A. Engelmann, S. A. Mihelic, A. Tomar, A. M. Hassan, and A. K. Dunn, “Evaluation of resonant scanning as a high-speed imaging technique for two-photon imaging of cortical vasculature,” Biomedical Optics Express 13(3), 1374–1385 (2022).

23. S. A. Mihelic, W. A. Sikora, A. M. Hassan, M. R. Williamson, T. A. Jones, and A. K. Dunn. “Segmentation-less, automated, vascular vectorization,” PLoS computational biology 17(10), e1009451 (2021).

24. B. Li, C. Wu, M. Wang, K. Charan, and C. Xu, “An adaptive excitation source for high-speed multiphoton microscopy,” Nature methods 17(2), 163–166 (2020).

