## supplemental figures for "Pulse train gating to improve signal generation for *in vivo* two-photon fluorescence microscopy"

### Figure S1

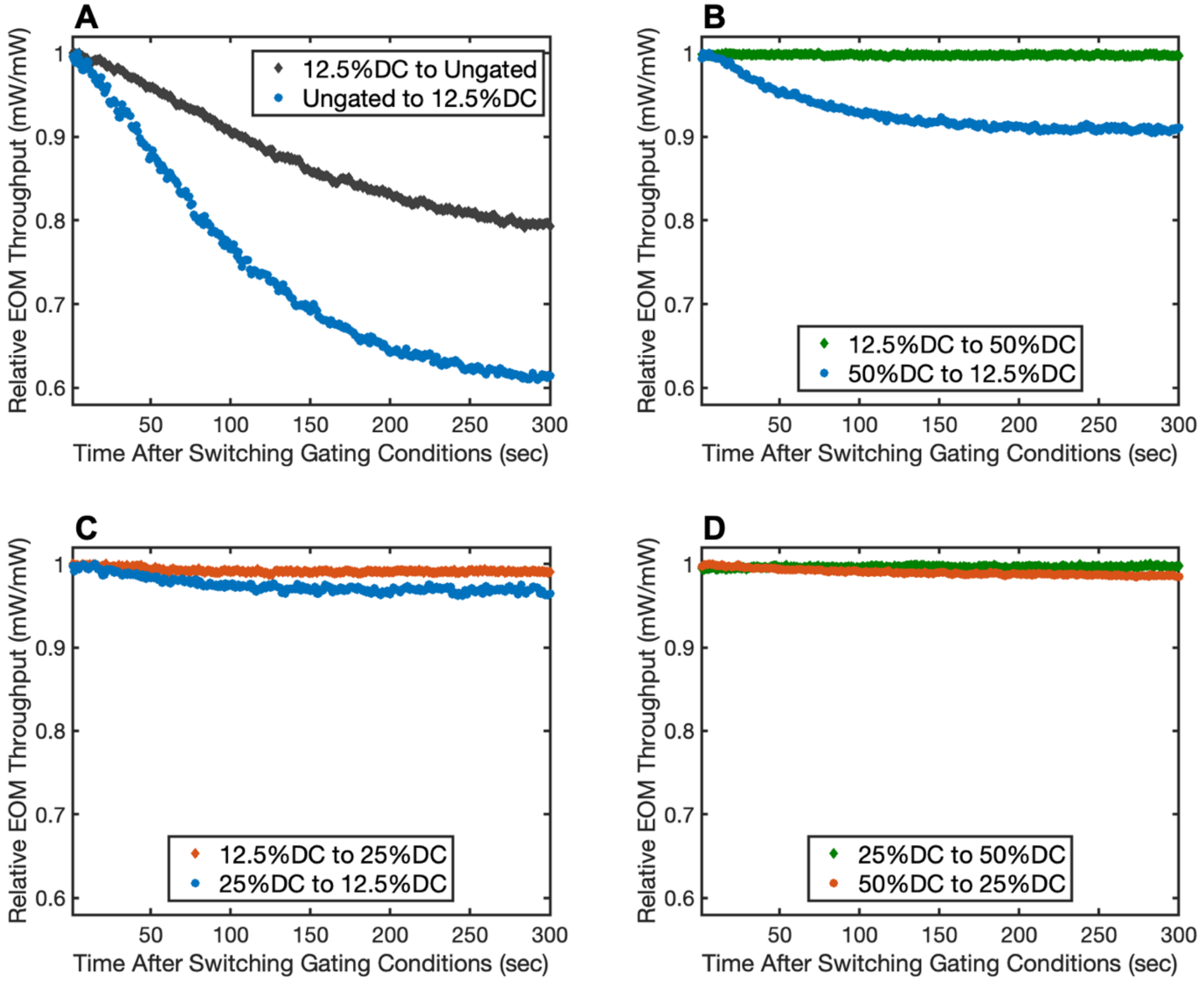


**Fig. S1** EOM throughput with time when switching gating conditions. (A) Switching from a 12.5% duty cycle to an ungated condition and vice versa. (B) Switching from a 12.5% duty cycle to a 50% duty cycle and vice versa. (C) Switching from a 12.5% duty cycle to a 25% duty cycle and vice versa. (D) Switching from a 25% duty cycle to a 50% duty cycle and vice versa.

### Figures S2 and S3

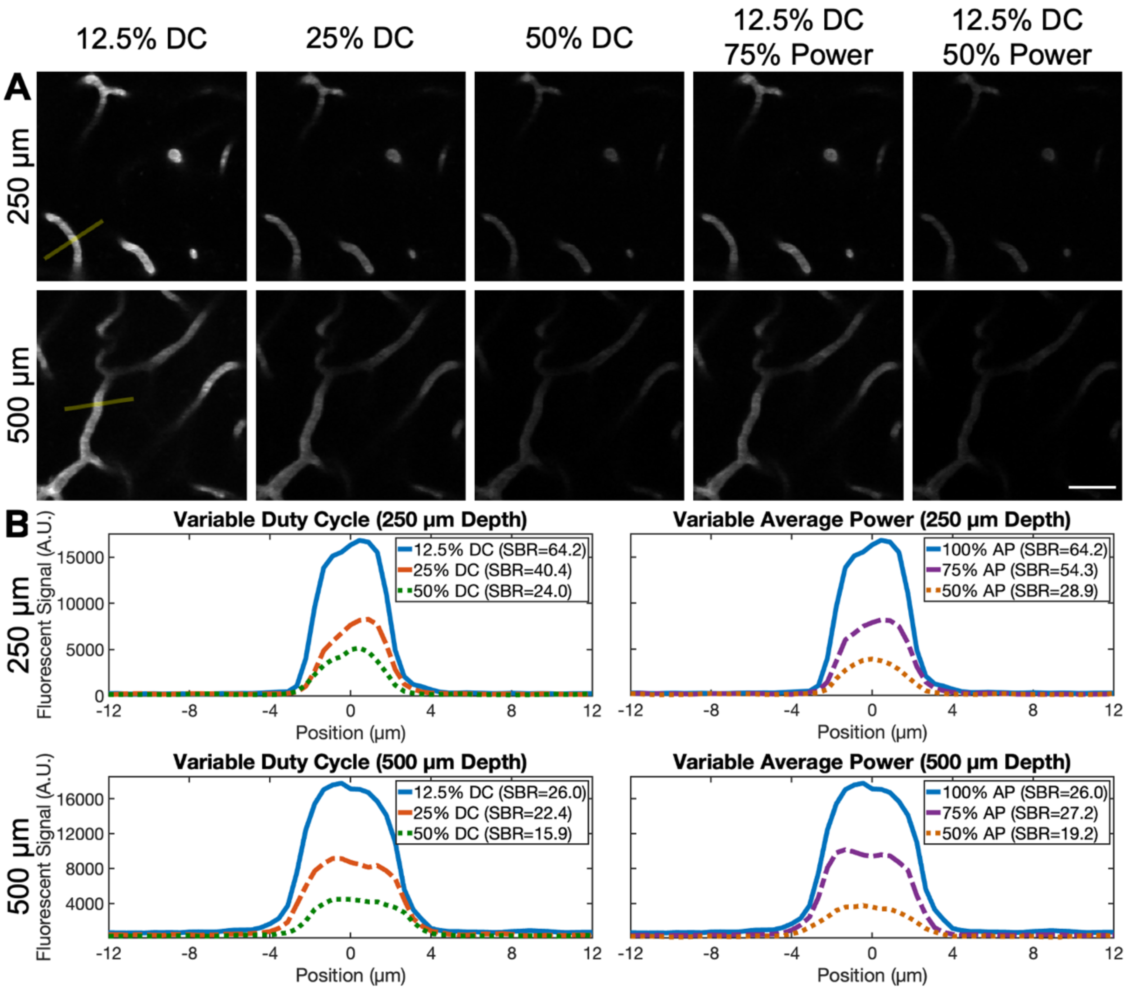


**Fig. S2** Second group of vessels (A) and line profiles (B) for Sec. 3.2.

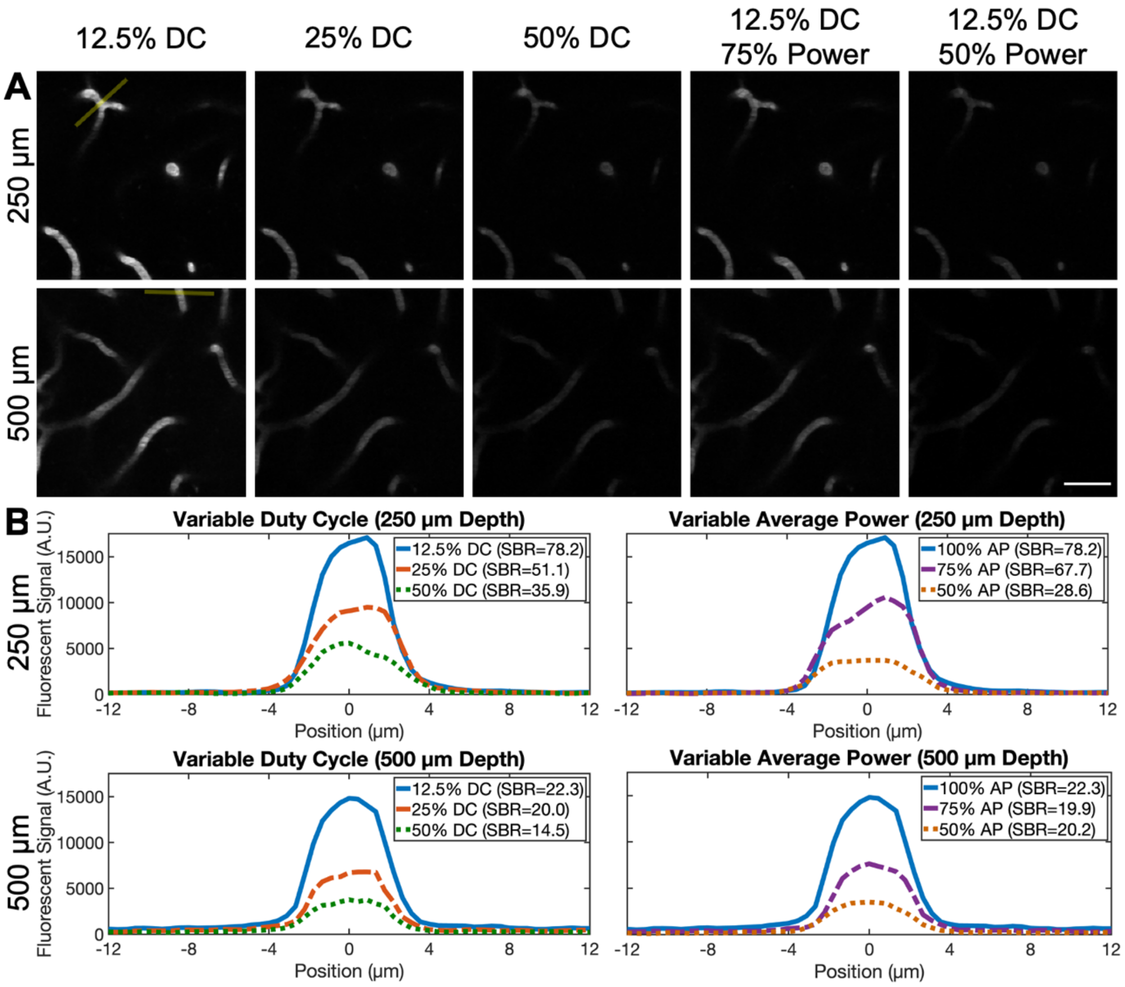


**Fig. S3** Third group of vessels (A) and line profiles (B) for Sec. 3.2

**Figure S4**

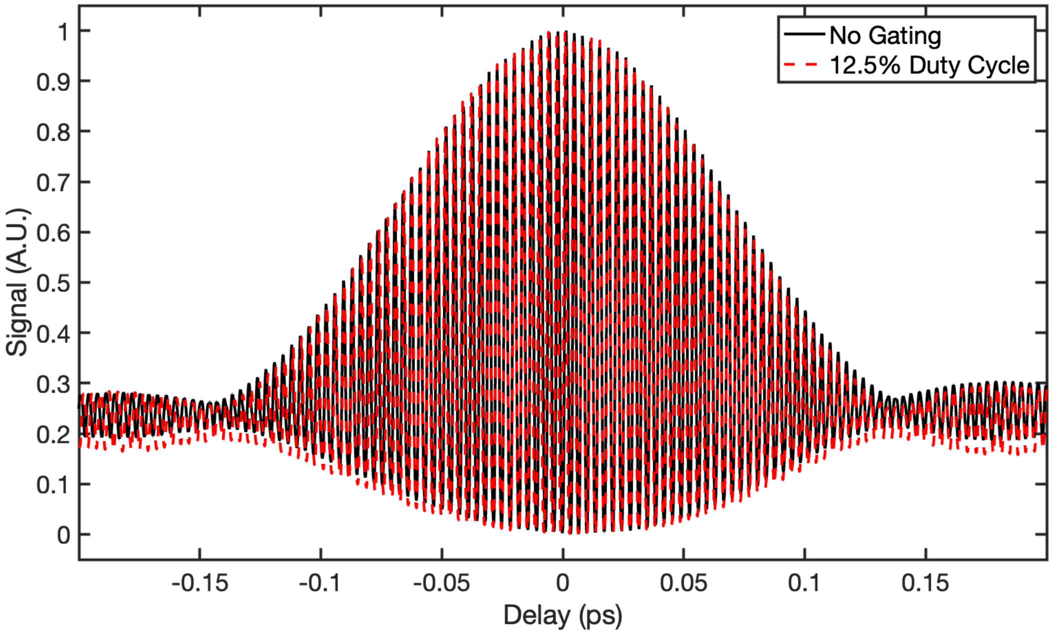


**Fig. S4** Comparison of autocorrelations for the ungated and 12.5% duty cycle excitation pulse trains. Autocorrelations were recorded using a tabletop autocorrelator (APE, pulseCheck).

.

### Figure S5

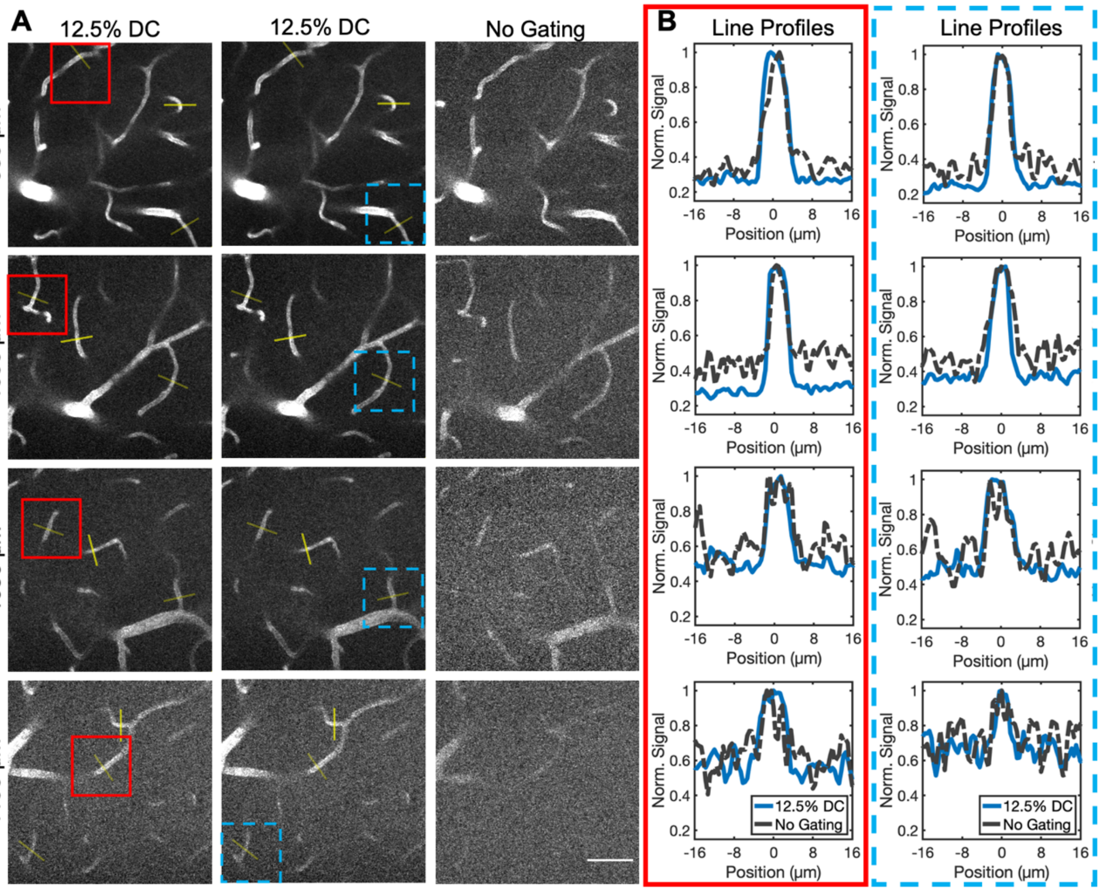


**Fig. S5** Images (A) and additional line profiles (B) that go with Sec. 3.3.
